# WaveMiner: a toolbox for navigating and analyzing the spatiotemporal properties of retinal waves

**DOI:** 10.1101/2025.10.23.684204

**Authors:** Kaylee M. Odum, Rishikesh K. Gupta, Elysia A. Gauthier, Alexander J. Fisch, Natalie A. Clark, Fernanda S. Orsi, Alexandre Tiriac

**Affiliations:** Department of Biological Sciences, Vanderbilt University, Nashville, TN 37232, USA; Department of Pharmacology, Vanderbilt University, Nashville, TN 37232, USA; Vanderbilt Brain Institute, Vanderbilt University, Nashville, TN 37232, USA; Department of Ophthalmology and Vision Sciences, Vanderbilt University, Nashville, TN 37232, USA

## Abstract

The precise development of the visual system is driven by retinal waves, which are bursts of spontaneous activity that propagate across retinal neurons in a wave-like fashion. In mice, retinal waves begin embryonically and continue until eye opening at the end of the second postnatal week. During this time, the mechanisms for generating and propagating retinal waves are changing, thus causing retinal waves to exhibit highly dynamic spatiotemporal properties from one day to the next. Critically, the spatiotemporal properties of retinal waves have been shown to instruct the development of the visual system, including eye-specific segregation, retinotopic mapping, direction selectivity, and potentially retinal vascularization. Currently, there is no method for the automatic detection and high-throughput quantitative analysis of the spatiotemporal properties of retinal waves. To overcome this barrier, we developed WaveMiner, an automated, high-throughput toolbox for detecting, segregating, and quantifying retinal waves from microelectrode-array (MEA) and calcium-imaging recordings. After first validating our toolbox, we use it to uncover novel dynamic spatiotemporal properties of retinal waves in the first two postnatal weeks. We also use this toolbox to analyze ultra long physiological recordings, revealing that waves exhibit both stable and dynamic spatiotemporal properties on an hourly basis. Finally, we demonstrate that this toolbox can detect waves in the presence of pharmacological agents that increase the baseline firing of neurons, enabling the discovery of novel factors that perturb the spatiotemporal properties of retinal waves and visual development. In summary, WaveMiner is a platform to standardize the detection and quantification of retinal waves across recording modalities and experimental conditions, enabling novel discoveries about their dynamic spatiotemporal properties and the factors that govern them.

**One Sentence Summary:** We present a novel toolbox for the automatic detection and analysis of retinal waves, which accurately captures their spatiotemporal dynamics across development and reveals new insights.

## Introduction

During early development of the vertebrate visual system before the onset of vision, the retina generates waves of spontaneous activity that propagate across the retinal ganglion cell (RGC) layer^1–4^. These retinal waves are critical for refining topographic maps, eye-specific segregation, and the emergence of feature selectivity in retinorecipient regions, such as the dorsal lateral geniculate nucleus (dLGN), superior colliculus (SC), and primary visual cortex^5–10^. In mice, retinal waves emerge in three distinct stages that correspond to the dynamic development of retinal circuits^11^. Stage I waves, occurring from embryonic day 16 (E16) through birth, are mediated by gap junctions and nicotinic acetylcholine receptors (nAChRs), and display diffuse, spatially variable activation^12–14^. Stage II waves, dominant during the first postnatal week (P1– P10), are driven by cholinergic signaling among starburst amacrine cells (SACs) via β2-containing nAChRs, and exhibit coherent, slowly propagating activity across large retinal territories^15,16^. Stage III waves, emerging around P10, are driven by glutamatergic signaling from bipolar cells, propagate more rapidly than Stage II waves, and activate ON and OFF RGCs sequentially^17,18^. Because the retinal circuitry changes continuously during development, wave properties differ measurably from one postnatal day to the next^19^.

Critically, retinal waves exhibit structured spatiotemporal properties that contribute meaningfully to circuit development^8^. These include defined wave frequencies (e.g.: approximately once per minute in stage II), spatial extent, and speeds that increase with age. One striking feature is the emergence of directional propagation, particularly a temporal-to-nasal bias during a transient window (P9–P12), which mimics optic flow experienced during forward locomotion^20–23^. This directionality has been shown to prime the development of motion-sensitive circuits in the SC^20^ and shape the dendrites of SACs^24^. Other spatiotemporal properties also matter: the slow frequency of stage II waves increases the likelihood of asynchronous activity between the two eyes which is thought critical for the development of eye-specific segregation^25^ and the area of waves matters for retinotopic mapping in binocular regions of the SC^26^. Lastly, the very act of a propagating wave leads to a local synchrony of neurons that is critical for Hebbian plasticity refinement of retinal axons in the SC^27^. The development of retinal vasculature is also thought to depend on retinal waves^28,29^, but whether this is done in an instructive or permissive manner is unknown. Across all stages, the patterned activity of retinal waves reflects the organization of the underlying circuitry and provides instructive signals for synaptic refinement and visual map formation before the onset of sensory experience. Accordingly, current questions in the field, like which circuits establish directional bias and whether models of neurodevelopmental disorders show altered wave structure^30,31^, are quantitative questions about retinal wave spatiotemporal properties, and progress depends directly on how reliably those properties can be measured.

Despite the recognized importance of retinal waves and their spatiotemporal dynamics, there is currently no widely adopted toolbox for automatic detection and characterization of retinal waves. Waves are three-dimensional objects extending in space (x, y) and time (t), and there is no agreed operational definition of a wave’s boundary, so metrics are not directly comparable across studies. Moreover, defining discrete wave events in noisy or complex datasets where tonic firing rates of RGCs change, whether from the more active stage III retinal waves and under pharmacological manipulations, remains a technical hurdle. Finally, prolonged recordings can contain thousands of waves, a volume that is impractical to curate by hand. Existing approaches are typically modality-specific (electrode arrays or calcium imaging), often rely on manual curation, and are based on custom algorithms that are not easily accessible. These limitations pose challenges for studying how wave properties evolve over time, respond to pharmacological and environmental perturbations, or vary across genetic backgrounds.

In this study, we introduce a new analytical toolbox designed to automatically segment and quantify retinal waves from data obtained using high-density microelectrode arrays (HD-MEA) and calcium imaging. We recorded a new dataset of retinal waves from birth to eye-opening in mice and used this to validate that our toolbox robustly captures known spatiotemporal features of waves across developmental stages. We discovered that at certain ages, a minority of waves contribute the bulk of activity, and establish a novel spatiotemporal metric that is critical for the refinement of retinal axons. We further demonstrate the toolbox’s utility in long recordings and under conditions of neural perturbation, providing novel insights about retinal waves. We also created a user-friendly application that utilizes all the tools present in our toolbox, which facilitates navigation and quality control of datasets. By standardizing wave detection and enabling new kinds of analyses, this toolbox aims to promote reproducibility and accelerate discovery of novel factors that govern retinal waves and visual system development.

## Results

### A toolbox to detect retinal waves and analyze their spatiotemporal properties

We developed a toolbox, called “WaveMiner”, capable of analyzing the comprehensive spatiotemporal dynamics of spontaneous retinal waves from *ex vivo* HD-MEA recordings of retinal activity^32^ (**figure 1A**). The toolbox loads data from virtually any retinal wave experiment that is structured in a way that contains both spatial and temporal information about neural activity, including HD-MEA electrophysiological data (**figure 1-5**) and calcium imaging data (**figure 6**). An initial module of the toolbox spatially aligns data across retinal wave experiments (temporal retina is always on the left, ventral retina is always on the bottom, and optic nerve is always at the center) allowing for simple and accurate regional spatiotemporal comparisons across recordings (**figure 1B**).

**Figure 1:**
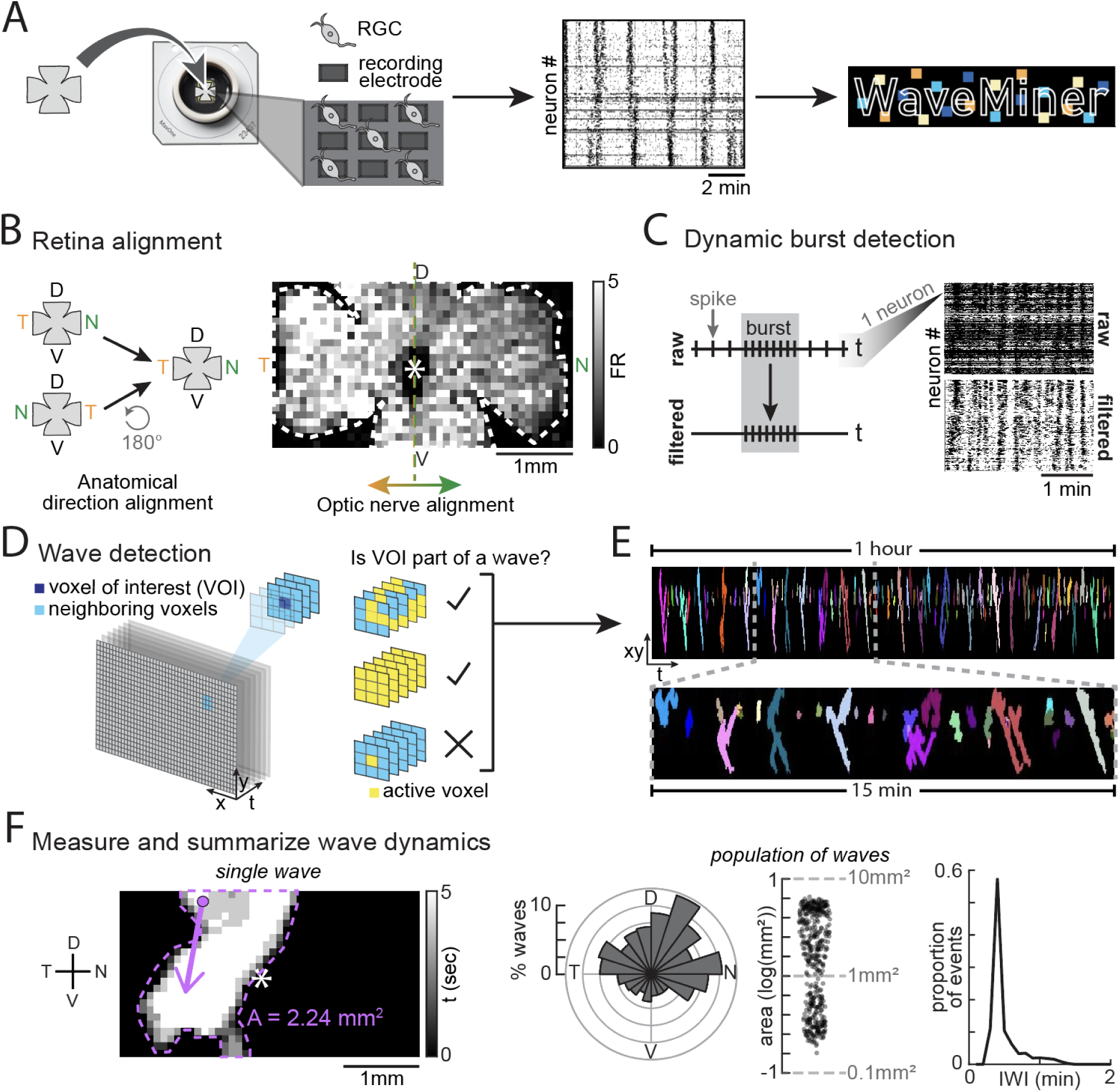
A toolbox to detect retinal waves and analyze their spatiotemporal properties. A) Spontaneous activity is captured by mounting the retina on a high-density microelectrode array (HD-MEA) chip so that the RGCs are touching the electrodes (left). The HD-MEA can record action potentials (spikes) from ∼1012 neurons simultaneously, as demonstrated by the raster plot of the recorded spikes from a P8 retina (middle). The recorded data can then be uploaded to the WaveMiner toolbox for wave segregation and analysis (right). B) Depiction of the toolbox’s retina alignment feature (left). The heatmap (right) shows the average spikes/sec (firing rate or FR) of each electrode over time in a recording of a P12 retina. The optic nerve is marked by the asterisk. C) The toolbox’s burst detection analysis filters tonic activity from the raw data, keeping only phasic activity (“bursts”). D) Schematic demonstrating how the toolbox detects and segregates retinal waves. When the algorithm finds an active voxel, it counts how many of its neighbors are active and determines whether the active voxel (VOI), is part of a wave. E) Representative image of segregated retinal waves after feeding data through the detection algorithm. Each color represents a different wave ID. F) Representation of the various spatiotemporal properties the toolbox can analyze both at the single wave level (left) and the wave population level (right, n=435 waves, N=1 P12 retina). Abbreviations: RGC – retinal ganglion cells, D – dorsal, N – nasal, V – ventral, T – temporal, FR – firing rate, VOI – voxel of interest, IWI – inter-wave interval

Before detecting the waves, the toolbox pre-processes the data by running a burst detection analysis to filter out tonic activity and keep phasic activity (which includes both propagating waves and transient, non-propagating events) (**figure 1C**). The burst detection filter is dynamic in time which allows for accurate filtering of phasic activity even across very long recordings where the tonic firing rate of neurons might change. The purpose of this phasic activity filtering is to allow the toolbox to detect waves in different conditions, especially ones where waves are present in conjunction with high tonic firing rates, like stage III waves or with pharmacological agents.

Once only phasic activity remains, the toolbox applies a 3D flood filling algorithm that automatically detects retinal waves. The algorithm is anchored on the definition that a unique wave constitutes any cluster of voxels connected in time (*t*) and space (*x* and *y*). To detect waves according to this definition, the algorithm scans every voxel of the xyt data matrix containing only phasic neural activity. Upon finding a voxel with activity, the algorithm begins segregating a wave from that voxel of interest (VOI) and analyzes neighboring voxels for activity (**figure 1D**). The process repeats until all active voxels connected in time and space are collected, forming a distinct retinal wave (more detailed information about this algorithm in **methods** section). Our algorithm successfully and reliably segregates retinal waves over hours of recordings, even in the presence of slow shifting baselines of tonic firing rates (**figure 1E**, **movie S1**, and **movie S2**). Following wave detection, the toolbox enables analysis of the spatiotemporal properties of individual waves (**figure 1F, left**) or populations of waves (**figure 1F, right**).

### Quantitative characterization of retinal waves across development using WaveMiner

To assess the accuracy of WaveMiner, we detected waves in retinal HD-MEA recordings taken at different ages and analyzed their spatiotemporal properties, which are well described (**figure 2A**). We binned the recordings into 5 age groups – 0 weeks (postnatal day (P)0-2, n=1182 waves, N=5 retinas across 5 mice), 0.5 weeks (P3-P5, n=575 waves, N=5 retinas across 5 mice), 1 week (P6-P8, n=691 waves, N=5 retinas across 4 mice), 1.5 weeks (P9-P11, n=1471 waves, N=5 retinas across 5 mice), and 2 weeks (P12-P14, n=1950 waves, N=5 retinas across 5 mice).

**Figure 2:**
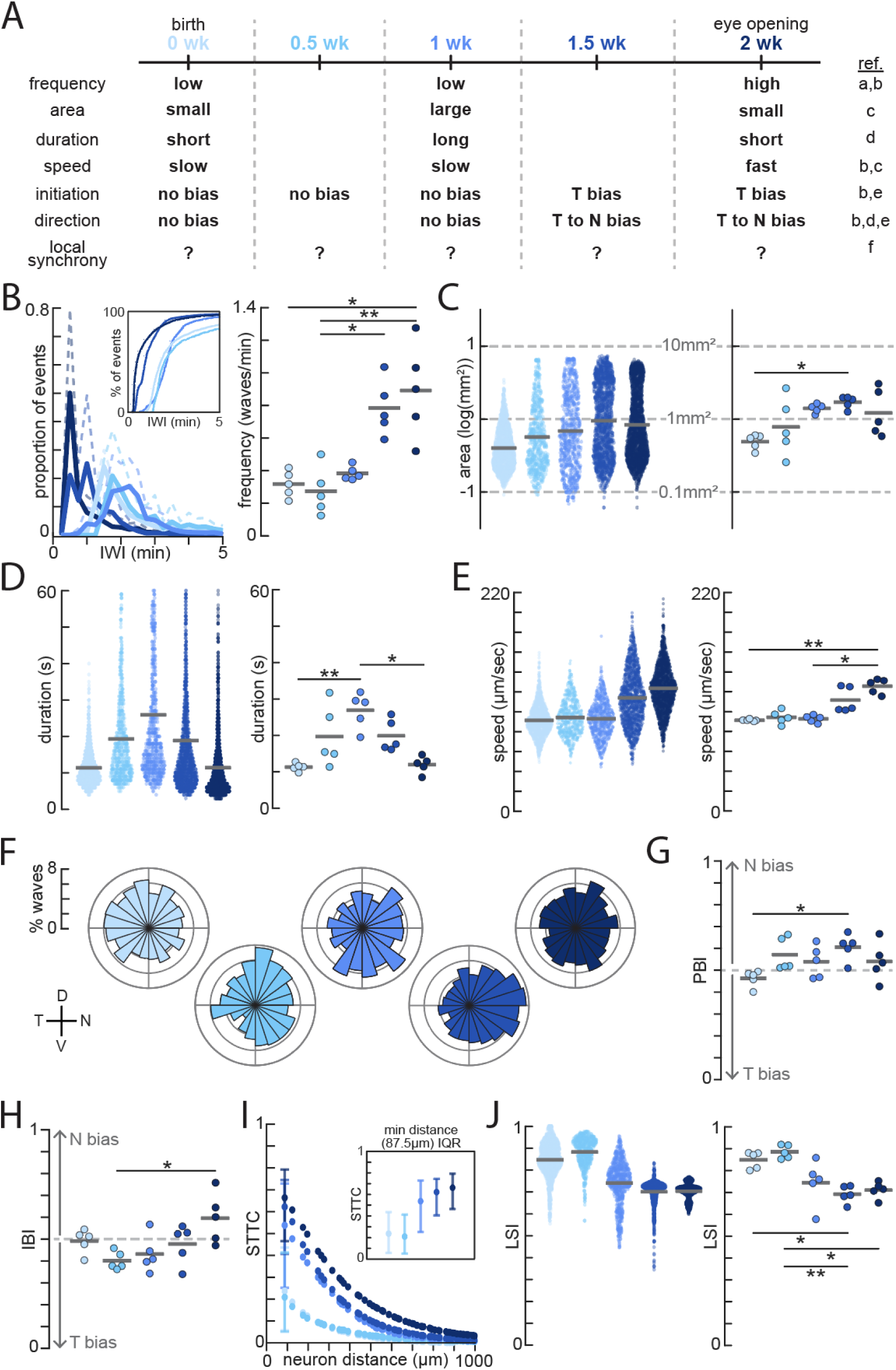
Quantitative characterization of retinal waves across development using WaveMiner. A) Developmental timeline of the spatiotemporal properties of retinal waves in mice. Age groups are defined as 0 weeks (P0-P2; n=1182 waves, N=5 retinas), 0.5 weeks (P3-P5; n=575 waves, N=5 retinas), 1 week (P6-P8; n=691 waves, N=5 retinas), 1.5 weeks (P9-P11; n=1471 waves, N=5 retinas), and 2 weeks (P12-P14; n=1950 waves, N=5 retinas). References: a – Cutts & Eglen, 2014^33^; b – Ackman et al., 2012^22^; c – Maccione et al., 2014^19^; d – Stafford et al., 2009^21^; e – Ge et al., 2021^20^; f – Matsumoto et al., 2024^27^. B) Left: Inter-wave intervals across age groups, shown as the proportion of events at a given interval length and as a cumulative distribution function. Solid lines represent the age group average and dotted lines represent the standard deviation. Right: Frequency of waves per mouse across age groups. \**p* < 0.05, \*\**p* < 0.01, \*\*\**p* < 0.005; Kruskal-Wallis followed by Dunn’s-Šidák post hoc test. C-E) Left: Area (shown on a log10 scale), duration, and top speed of individual waves across age groups. These are descriptive, non-inferential data. Right: Average wave area (shown on a log10 scale), duration, and top speed per retina across age groups. \**p* < 0.05, \*\**p* < 0.01, \*\*\**p* < 0.005; Kruskal-Wallis followed by Dunn’s-Šidák post hoc test. F) Polar histograms depicting propagation direction of waves across ages groups. G) Propagation Bias: Proportion of waves per retina that propagated in the nasal direction (PBI), analyzed on one axis (temporal vs. nasal direction). The dashed line at PBI=0.5 indicates no bias. \**p* < 0.05, \*\**p* < 0.01, \*\*\**p* < 0.005; Kruskal-Wallis followed by Dunn’s-Šidák post hoc test. H) Initiation Bias: Proportion of nasally initiated waves (IBI) per retina across ages groups, analyzed by halves (temporal vs. nasal retina). The dashed line at IBI=0.5 indicates no bias. \**p* < 0.05, \*\**p* < 0.01, \*\*\**p* < 0.005; Kruskal-Wallis followed by Dunn’s-Šidák post hoc test. I) Median STTC at each unique neuron distance across age groups (Δt=0.5; 0wk, n=380854 neuron pairs, N=5 retinas; 0.5wk, n=341299 neuron pairs, N=5 retinas; 1wk, n=578017 neuron pairs, N=5 retinas; 1.5wk, n=649140 neuron pairs, N=5 retinas; 2wk, n=487031 neuron pairs, N=5 retinas). The interquartile range (IQR) is only shown at the minimum distance for graph readability. At the minimum neuron distance (87.5μm), 0wk median=0.237 (IQR [0.057-0.436]; n=6073 neuron pairs, N=5 retinas), 0.5wk median=0.209 (IQR [0.051-0.413]; n=5335 neuron pairs, N=5 retinas), 1wk median=0.537 (IQR [0.252-0.729]; n=7843 neuron pairs, N=5 retinas), 1.5wk median=0.622 (IQR [0.407-0.745]; n=8721 neuron pairs, N=5 retinas), 2wk median=0.663 (IQR [0.465-0.792]; n=7029 neuron pairs, N=5 retinas). J) Left: Local synchrony index (LSI) of individual waves across age groups, calculated as the ratio of activity ahead of wavefront versus activity within wavefront. These are descriptive, non-inferential data and statistical tests were run on the data shown on the right. Right: Average LSI per retina across age groups. \**p* < 0.05, \*\**p* < 0.01, \*\*\**p* < 0.005; Kruskal-Wallis followed by Dunn’s-Šidák post hoc test. Abbreviations: wk – week, IWI – inter-wave interval, D – dorsal, N – nasal, V – ventral, T – temporal, PBI – propagation bias index, IBI – initiation bias index, STTC – spike timing tiling coefficient, LSI – local synchrony index.

We first assessed the occurrence of waves at the various age groups, both in terms of inter-wave-intervals (IWIs) (**figure 2B, left**), which calculates the interval of time between the onset of waves per unit space of the retina, and frequency (**figure 2B, right**), which represents the occurrence of waves per minute per unit space of the retina. Both analyses revealed that waves are much more frequent in the 2^nd^ postnatal week than the 1^st^. We next assessed the areas of waves and found that they gradually become larger throughout development before shrinking back in late stage III (**figure 2C**). When assessing the duration of waves, we observed that waves exhibit a gradual increase peaking at 1 week, before gradually decreasing in the 2^nd^ week (**figure 2D**). Lastly, analyzing speed revealed waves were much faster in the 2^nd^ postnatal week than the 1^st^ (**figure 2E and figure S2**). Overall, these results are consistent with previous reports^19–22,27,33^(**figure 2A**).

We next tested whether waves exhibit any location-specific biases during the first two postnatal weeks, both regarding propagation and initiation bias. We first assessed the propagation direction of retinal waves in each age group. Retinas from 1.5-week-old mice exhibited a markedly temporal-to-nasal propagation bias (**figure 2F-G**), consistent with both *ex vivo*^21,34^ and *in vivo*^20^ reports. We next assessed the initiation location of retinal waves in each age group. Retinas from 0.5-1-week-old mice exhibited a slight initiation bias in temporal retina (**figure 2H**), consistent with *in vivo* reports^22^.

Lastly, we examined the signal-to-noise characteristics of retinal activity during development in both a wave-independent and wave-dependent manner. We first calculated the spike time tiling coefficient (STTC), a measure of the correlation between pairs of neurons that is independent of firing rate^33^. Because STTC was calculated on raw spiking data (prior to wave segregation), this analysis is wave independent. Previously, STTC was calculated on retinal wave HD-MEA recordings from mouse retinas at P9, P15, P21, and 6 weeks, and it was reported that STTC decreases with age^33^. However, we found that STTC of nearby neurons began low (∼0.2-0.3) at 0 weeks and gradually increased until it reached its highest (∼0.6-0.7) at 2 weeks (**figure 2I**). The STTC of our P9-P11 age group was consistent with the values reported previously^33^. We also assessed local synchrony, a wave-dependent measure of how much activity is at the wavefront versus just ahead of the wavefront and a feature of retinal waves thought to be important for the refinement of RGC axons^27^. We found that local synchrony was strongest during early development and gradually became weaker over time (**figure 2J**).

### A minority of waves accounts for the majority of retinal activity after the first postnatal week

The combination of HD-MEAs, large recordings of the retinal surface area, and automatic detection provided by WaveMiner results in a large dataset of retinal waves: 5869 waves across 25 retinas (24 mice). The scale of this dataset revealed that although large retinal waves emerge in the second postnatal week (**figure 2C**), small retinal waves remain throughout development. As a consequence, the range of wave sizes widened with development, with the second postnatal week containing both small and large waves.

This distribution has practical implications: A small, transient wave and a large, lingering wave are counted equally in population averages, yet they contribute unequally to the overall activity a developing retina actually experiences. Thus, any assessment of genetic or pharmacological manipulation on retinal waves could be greatly affected by an assessment that counts waves equally. To quantify whether this is the case, we assigned each wave an activity impact factor (AIF), defined as the number of voxels that wave activated as a fraction of all voxels activated by waves in the recording, essentially computing the proportion of activity a wave contributed during that recording. Ranking waves by AIF and taking the cumulative sum then describes how activity impact is distributed across the population of retinal waves. This distribution changed gradually with age (**figure 3A** and **figure S1**). We calculated the Gini coefficient for each retina, which measures how close a curve is to absolute equality (a perfectly linear line). A value of 0 indicates total equality, while a value of 1 indicates total inequality.

**Figure 3:**
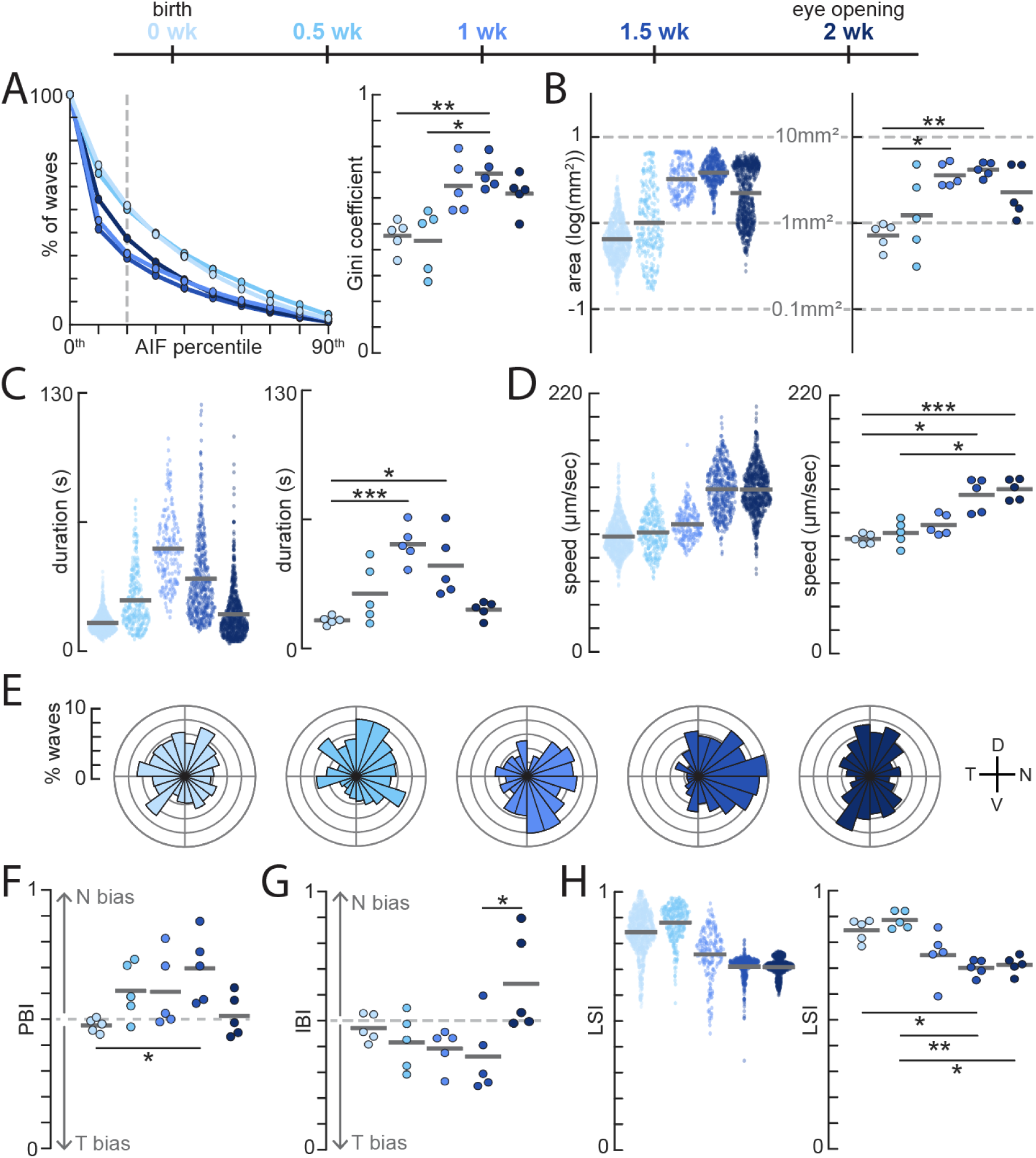
A minority of waves accounts for the majority of retinal activity after the first postnatal week. A) Left: Percentage of waves within each activity impact factor (AIF) percentile per retina across age groups. Each thin line represents data from one retina the thick lines represent the average for the age group. Only waves that fall within the 20^th^ AIF percentile (including and to the right of the dashed line) were analyzed in B-H (0wk, n=785 waves, N=5 retinas; 0.5wk, n=281 waves, N=5 retinas; 1wk, n=197 waves, N=5 retinas; 1.5wk, n=436 waves, N=5 retinas; 2wk, n=750 waves, N=5 retinas). Right: Quantification of the data on the left. A Gini coefficient was calculated for each retina across age groups based on its Lorenz curve. A Gini coefficient of 1 indicates all waves contributed equally. \**p* < 0.05, \*\**p* < 0.01, \*\*\**p* < 0.005; Kruskal-Wallis followed by Dunn’s-Šidák post hoc test. B-D) Left: Area (shown on a log10 scale), duration, and top speed of individual waves in the 20^th^ AIF percentile across age groups. These are descriptive, non-inferential data. Right: Average wave area (shown on a log10 scale), duration, and top speed of waves in the 20^th^ AIF percentile per retina across age groups. \**p* < 0.05, \*\**p* < 0.01, \*\*\**p* < 0.005; Kruskal-Wallis followed by Dunn’s-Šidák post hoc test. E) Polar histograms depicting propagation direction of waves in the 20^th^ AIF percentile across ages groups. F) Propagation Bias: Proportion of waves in the 20^th^ AIF percentile per retina that propagated in the nasal direction (PBI), analyzed on one axis (temporal vs. nasal direction). The dashed line at PBI=0.5 indicates no bias. \**p* < 0.05, \*\**p* < 0.01, \*\*\**p* < 0.005; Kruskal-Wallis followed by Dunn’s-Šidák post hoc test. G) Initiation Bias: Proportion of nasally initiated waves (IBI) in the 20^th^ AIF percentile per retina across ages groups, analyzed by halves (temporal vs. nasal retina). The dashed line at IBI=0.5 indicates no bias. \**p* < 0.05, \*\**p* < 0.01, \*\*\**p* < 0.005; Kruskal-Wallis followed by Dunn’s-Šidák post hoc test. H) Left: Local synchrony index (LSI) of individual waves in the 20^th^ AIF percentile across age groups. These are descriptive, non-inferential data. Right: Average LSI of waves in the 20^th^ AIF percentile per retina across age groups. \**p* < 0.05, \*\**p* < 0.01, \*\*\**p* < 0.005; Kruskal-Wallis followed by Dunn’s-Šidák post hoc test. Abbreviations: wk – week, AIF – activity impact factor, D – dorsal, N – nasal, V – ventral, T – temporal, PBI – propagation bias index, IBI – initiation bias index, LSI – local synchrony index.

At 0 and 0.5 weeks, waves contributed to total retinal activity relatively equally: population size declined near-linearly as the cumulative AIF threshold increased (**figure 3A, left**) and the Gini coefficient was relatively low (**figure 3A, right**). From 1 week onward, the decline in population size became progressively steeper and the Gini coefficient increased significantly, indicating that the overall phasic activity that retinas experienced were increasingly due to a minority of large, long-lasting waves. By 1-2 weeks, ∼40% of retinal waves accounted for 80% of total phasic activity. The overall impact of retinal activity therefore becomes increasingly concentrated to a subset of retinal waves across development.

The concentration of activity impact on a minority of waves matters for how wave properties are measured. When a minority of waves carries most of the activity that a retina experiences, population-wide averages of area, duration, and speed are weighted heavily by numerous low-impact events and may obscure changes in the waves that deliver most of the instructive signal. We therefore re-examined the developmental trajectory using only the waves that contribute 80% of the overall retinal activity (waves in the 20^th^ AIF percentile) per recording (**figure 3B-H**). Thus, narrowing the analysis to high-impact waves results in distributions of spatiotemporal properties that more closely match previously published values: primarily broad waves in the middle of stage II waves, a more pronounced jump in wave speeds at the start of stage III waves, and a sharper propagation bias at 1.5 weeks ^13,20,26,28–30^.

### Coupling WaveMiner with long HD-MEA recordings reveals the dynamic and stable nature of spatiotemporal properties of retinal waves

One outstanding question about retinal waves is whether the spatiotemporal properties remain stable or change dynamically over the course of several hours. To begin to answer this question using WaveMiner, we performed nine-hour long HD-MEA recordings of retinal waves in 0.5-week-old mice (P5: n=3039 waves, N=3 retinas across 3 mice) and in 1.5-week-old mice (P12: n=6892 waves, N=2 retinas across 2 mice) (**figure 4A, left**). Nine-hour recordings combined with the ability to sample across large areas of the retina enabled a high throughput assessment of the spatiotemporal properties of waves in individual mice (**figure 4A, right**).

**Figure 4:**
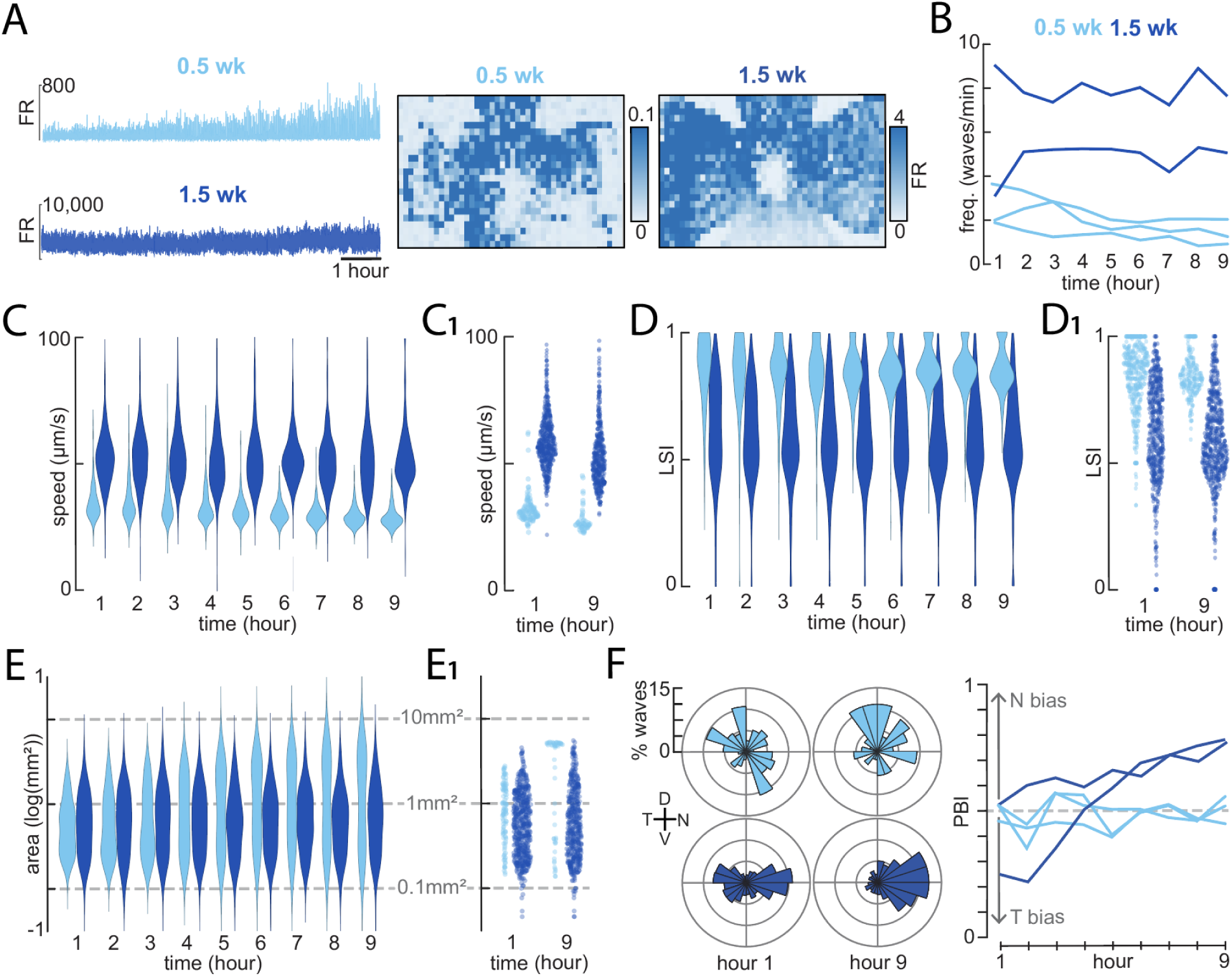
Long term recordings coupled with WaveMiner reveal the dynamic and stable structure of spatiotemporal properties of waves. A) Left: Representative retinal activity obtained from nine-hour long HD-MEA recordings, one from a 0.5-week-old mouse (light blue) and one from a 1.5-week-old mouse (dark blue). Right: Heatmaps of mean firing rate of neurons recorded over the course of nine hours. B) Wave frequency over the course of nine hours in 0.5-week-old (n=3039 waves, N=3 retinas) and 1.5-week-old (n=6892 waves, N=2 retinas) mice. Each line represents data from one retina. C) Average speed of individual waves recorded across all 0.5- and 1.5-week-old retinas, binned per hour. C1) Average speed of individual waves from a representative 0.5-week-old (n=685 waves) and 1.5-week-old (n=4309 waves) retina at the first hour and ninth hour of the recording. Data are from the same representative recordings as in (A). D) Local synchrony index (LSI) of individual waves recorded across all 0.5- and 1.5-week-old retinas, binned per hour. D1) Local synchrony index (LSI) of individual waves from a representative 0.5- and 1.5-week-old retina at the first hour and ninth hour of the recording. Data are from the same representative recordings as in (A). E) Wave area (on a log10 scale) of individual waves recorded across all 0.5- and 1.5-week-old retinas, binned per hour. E1) Wave area (on a log10 scale) of individual waves from a representative 0.5- and 1.5-week-old retina at the first hour and ninth hour of the recording. Data are from the same representative recordings as in (A). F) Left: Polar histograms depicting propagation direction of waves in the first and ninth hour for a representative 0.5- and 1.5-week-old mouse. Data are from the same representative recordings as in (A). Right: Propagation bias calculated per hour over the course of the nine hours. Each line represents data from one retina. Abbreviations: wk – week, FR – firing rate, LSI – local synchrony index, D – dorsal, N – nasal, V – ventral, T – temporal, PBI – propagation bias index.

We found several spatiotemporal properties that were consistently stable across the duration of the nine-hour recording despite exhibiting age-dependent properties. For example, 0.5-week-old retinas exhibited less frequent waves than 1.5-week-old retinas, but both had stable frequencies over the nine-hour recordings (**figure 4B**). Likewise, although wave speeds were slower for 0.5-week-old retinas than 1.5-week-old retinas, all wave speeds were stable throughout the duration of the recordings (**figure 4C and C1**). Similarly, although local synchrony was higher for 0.5-week-old retinas than 1.5-week-old retinas, all local synchrony measures were stable throughout the duration of the recordings (**figure 4D and D1**).

Surprisingly, we found a couple spatiotemporal properties that were dynamic for some ages but stable for others. First, in our recordings of 0.5-week-old retinas, we observed that the distribution of waves areas was unimodal in the first hour but bimodal by the ninth hour, with both very large and very small waves present (**figure 4E and E1**). In our recordings for 1.5-week-old retinas, the distribution of wave areas is stable across the nine hours. Second, in our recordings for 1.5-week-old retinas, the propagation of retinal waves did not initially exhibit a nasal-dominant bias but developed a strong nasal-dominant bias over the course of the nine hours (**figure 4F**). In our recordings for 0.5-week-old retinas, retinal waves exhibited stable symmetric propagation (no bias) across the nine hours. All together, these results demonstrate that the spatiotemporal properties of retinal waves exhibit distinct dynamic features depending on the age of the animal.

### WaveMiner enables the detection of retinal waves even in high noise environments

A key method to discover circuits and factors that govern retinal waves is to use pharmacology or genetic manipulations that perturb retinal waves. Unless the manipulation completely abolishes retinal waves, determining the effect of the manipulation on the spatiotemporal properties of retinal waves is difficult. Calculations that are agnostic to retinal waves, like the STTC calculation^33^, allow information about how a manipulation affects distance-dependent synchrony, but not how other spatiotemporal properties of waves, like area or propagation bias, are affected. Thus, we tested whether our toolbox could continue to reliably detect retinal waves even under the most severe cases of neural activity manipulation. To do so, we utilized mice where the excitatory Gq-DREADD is expressed in all RGCs (Vglut2-ires-cre::Gq-DREADD) and performed 10-minute HD-MEA recordings before and after activating DREADDs with clozapine-n-oxide (CNO). We furthermore performed these experiments in the 1^st^ and 2^nd^ postnatal week (PW) to test whether we could segregate cholinergic and glutamatergic waves in high noise conditions.

For both the first and second postnatal week, CNO applied on Gq-DREADD-expressing retinas elicited a large increase in the tonic firing rate of retinal neurons (**figure 5A-B**), thereby increasing the overall “noise” in the HD-MEA recording. Our toolbox still successfully segregated retinal waves despite this prominent increase in noise (**figure 5C** and **movie S3**) and found a comparable number of waves before and after application of CNO (PW1: pre-CNO n=228 waves, post-CNO n=261 waves, N=8 retinas across 6 mice; PW2: pre-CNO n=372 waves, post-CNO n=286 waves, N=6 retinas across 4 mice). We also assessed their spatiotemporal properties (**figure 5D-I**). Most notably, following activation of Gq-DREADDs, retinal waves propagated faster and exhibited a strong decrease in local synchrony.

**Figure 5:**
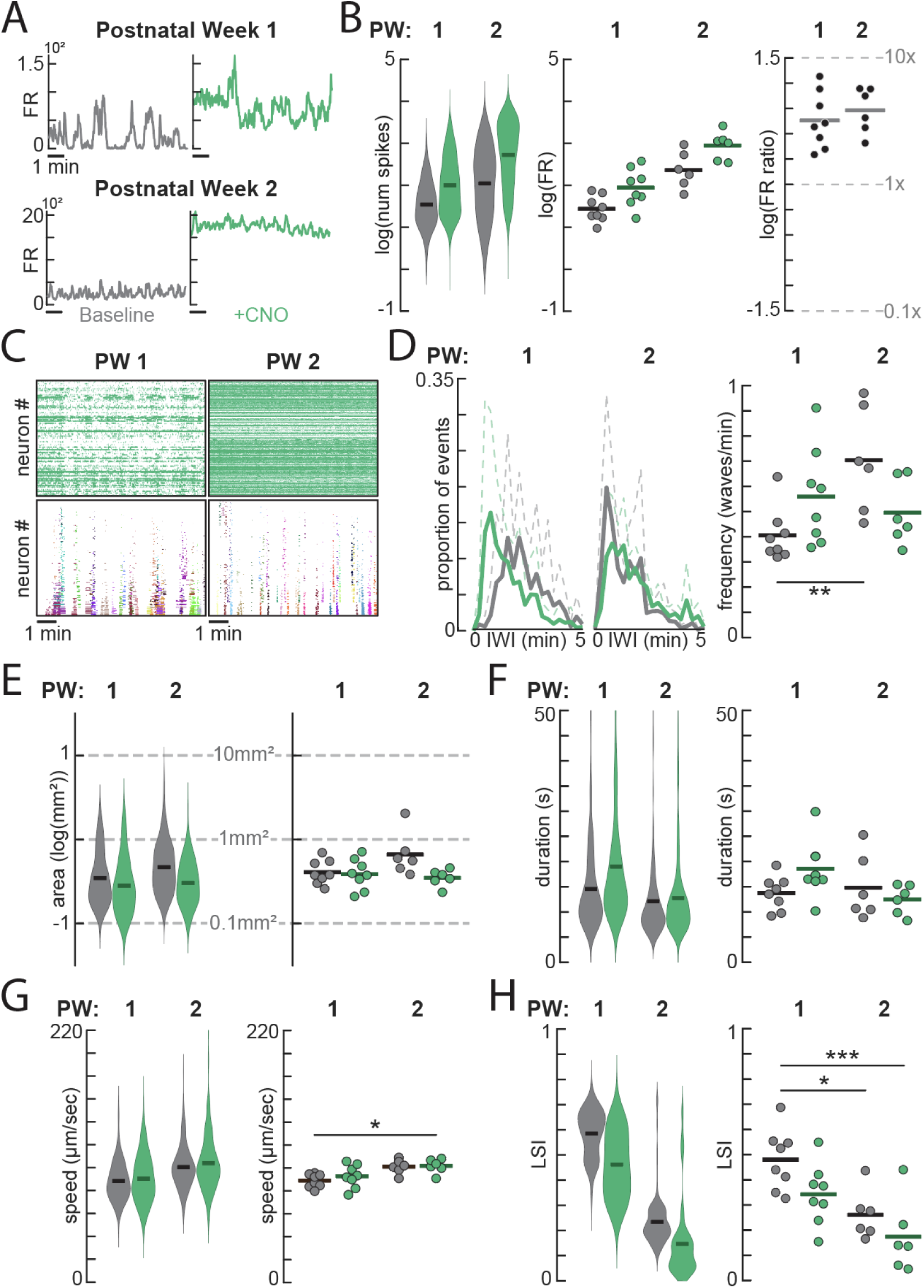
Using WaveMiner to segregate and analyze waves in recordings with very high noise. A) Representative data of the sum of spikes across the retina across a 10-minute HD-MEA recording, both before (black) and after (green) application of 10µM CNO on retinas from vGlut2-ires-cre::Gq-DREADD mice in the first postnatal week (top, P5, N=1 retina) and the second postnatal week (bottom, P12, N=1 retina). B) Left: Total number of spikes per neuron before (black) and after (green) application of CNO during the first postnatal week (PW1, n=2999 neurons, N=8 retinas) and the second postnatal week (PW2, n=3986 neurons, N= 6 retinas) on a log10 scale. Middle: Average firing rate or FR (spikes/sec) per retina across age groups before (black) and after (green) application of CNO. Right: FR ratio (post-CNO FR/pre-CNO FR) on a log10 scale per retina across age groups, showing the fold change in activity (PW1 average = 3.16x and PW2 average = 3.80x). C) Representative raster plots of recorded spikes per neuron following the application of CNO in vGlut2-cre/Gq-DREADD mice in the first postnatal week (top left, P5) and the second postnatal week (top right, P12) and the detected waves in each recording (bottom left, P5 and bottom right, P12). Each color represents a unique wave ID (P5, n=40 waves, N=1 retina; P12, n=42 waves, N=1 retina). D) Wave frequency per retina before (black) and after (green) application of CNO across age groups (PW1, N=8 retinas; PW2, N=6 retinas). E) Inter-wave-intervals (IWIs) between consecutive waves before (black) and after (green) application of CNO during PW1 and PW2, shown as a proportion of events at any given IWI. Dashed lines represent standard deviation. F-I) Left: Area (on a log10 scale), duration, top speed, and local synchrony (LSI) of individual waves before (black) and after (green) application of CNO across age groups (PW1: pre-CNO n=228 waves, post-CNO n=261 waves, N=8 retinas; PW2: pre-CNO n=372 waves, post-CNO n=286 waves, N=6 retinas. These are descriptive, non-inferential data and all statistical tests were run on the data shown on the right. Right: Average wave area (shown on a log10 scale), duration, and top speed per retina before (black) and after (green) application of CNO across age groups. \**p* < 0.05, \*\**p* < 0.01, \*\*\**p* < 0.005; Kruskal-Wallis followed by Dunn’s-Šidák post hoc test. Abbreviations: FR – firing rate, PW – postnatal week, LSI – local synchrony index

### Using WaveMiner to segregate and analyze waves in calcium imaging data

To make our toolbox more widely applicable, we adapted WaveMiner into a version that processes data from calcium imaging, another technique commonly used to capture retinal waves. The primary difference between the original HD-MEA version of WaveMiner and the calcium imaging version occurs prior to wave detection due to the nature of the raw data. HD-MEA data contains spikes, while calcium imaging data contains transients (**figure 6A, middle**). Thus, the pre-processing steps differ (see **methods** for more details), but the data format is essentially identical by the time it reaches the wave detection step of the toolbox (**figure 6A, right**). At this point, the calcium imaging version of WaveMiner detects waves using the same flood filling algorithm described previously (**figure 1D**) and segregates the waves into a 3D matrix, where each wave has a unique wave ID (**figure 6B** and **movie S4**). To further validate WaveMiner’s ability to process calcium imaging data, we analyzed several spatiotemporal properties of embryonic retinal waves using previously published data^12^. IWI, frequency, and area (n=623 waves, N=4 retinas) were all consistent with the values reported in the study^12^ (**figure 6C-D**). To further test the calcium imaging version of WaveMiner, we mapped the initiation sites and vector flow field of a representative recording (n=76 waves) (**figure 6E-F**).

**Figure 6:**
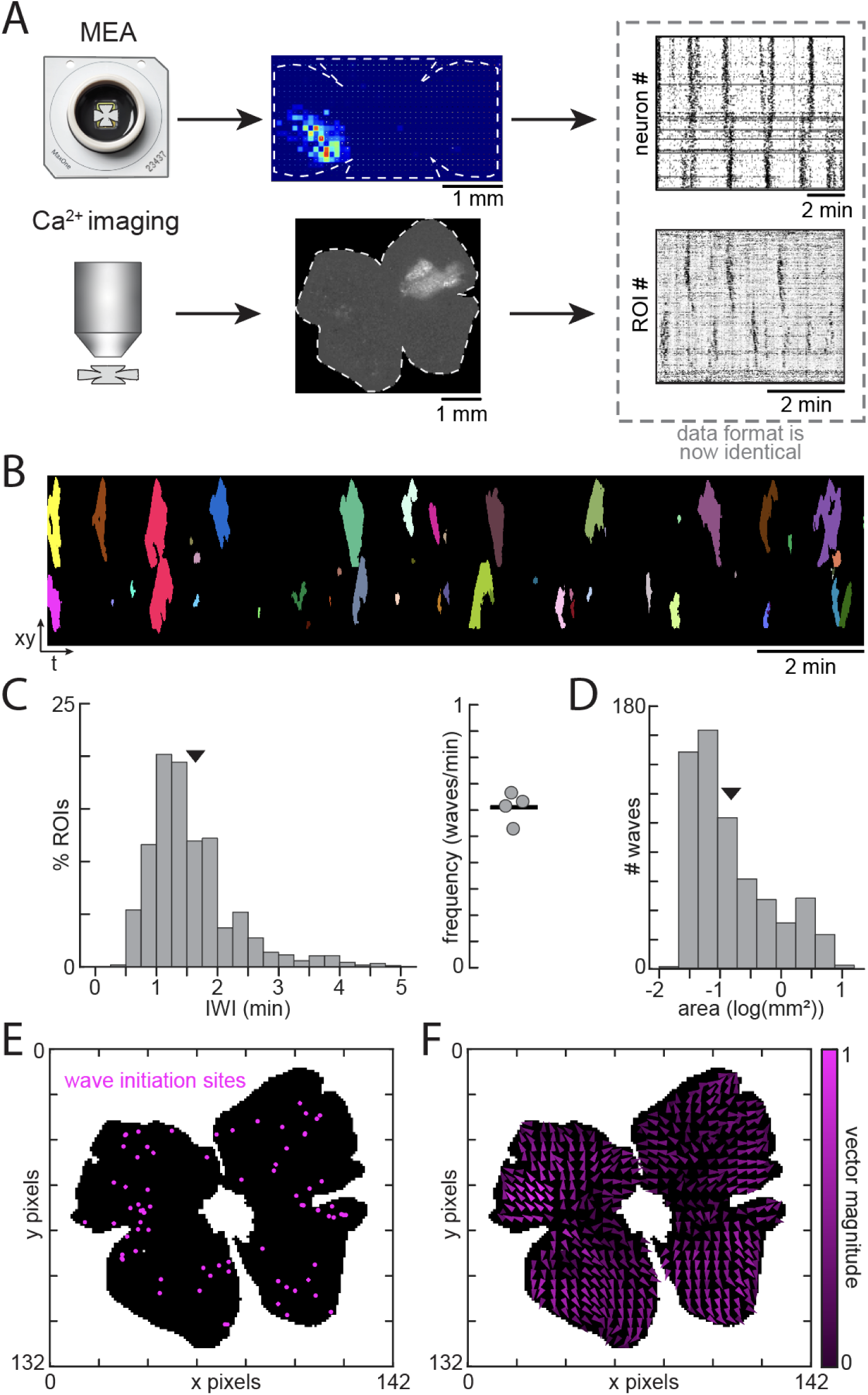
WaveMiner can segregate and analyze waves in Ca^2+^ imaging data. A) In addition to HD-MEA data, the toolbox can analyze Ca^2+^ imaging data (or any data in .tif format). While the initial data processing steps differ (see methods), the data format is essentially identical when it is fed into the wave detection algorithm. B) Representative image of waves segregated from the Ca^2+^ imaging recording of embryonic retinal waves shown in (A)^12^. Each color represents a different wave ID. C) Inter-wave intervals, shown as the percent of ROIs at a given interval length (left; n=623 waves, N=4 retinas) and the wave frequency per retina (right). Arrow represents the average IWI. D) Wave area, shown as the number of waves at a given area along a log10 distribution (n=623 waves, N=4 retinas). Arrow represents the average area. E) Initiation sites of waves from the example Ca^2+^ imaging recording in (A) (n=76 waves). F) Flow field showing the net flow of all waves from the example Ca^2+^ imaging recording in (A) (n = 76 waves). The vector magnitude refers to the number of waves that pass over a given pixel in the net direction. Abbreviations: ROI – region of interest, IWI – inter-wave interval

## Discussion

Here we report on a new tool that we built to segregate individual retinal waves and analyze their spatiotemporal properties. Our toolbox, WaveMiner, was designed to work on both HD-MEA and calcium imaging data but can work with any dataset that has spatial and temporal information. We validated our toolbox on the known spatiotemporal properties of retinal waves during development: namely our toolbox correctly catches that the increasing frequency and speed of waves that occurs in the second week, correctly identifies the propagation bias in the middle of the second postnatal week, and identifies the large waves seen during the stage II developmental period. The combination of using HD-MEAs, large surface area recordings of retinal waves, and automatic detection has revealed a novel phenotype about retinal waves, which is that the overall activity impact is heavily concentrated into a minority of retinal waves as mice age. In addition to calculating known spatiotemporal properties, we added novel analyses, like the local synchrony, that have become relevant given recent publications^27^. We used this toolbox to analyze the spatiotemporal properties of retinal waves in long recordings and demonstrated that spatiotemporal properties exhibit stable or dynamic nature in an age-dependent manner. We also demonstrated that our toolbox can detect and analyze retinal waves following the addition of pharmacological agents, enabling the assessment of how retinal waves are impacted by various manipulations that affect the tonic firing rate of RGCs. Finally, we demonstrated that our toolbox can be applied to calcium imaging data and validated it accordingly^12^. Our toolbox can be used by running a pipeline of codes that are available on our GitHub page or by running a standalone graphical user interface (GUI). In summary, our toolbox has broad applicability, has been extensively validated with known results from the literature, will help standardize results across labs and methodologies, and is designed to be user friendly.

### The analytical impact of the increased concentration of activity impact over development

Our results show that as mice age, the overall amount of phasic activity that the retina experiences is attributed to a minority of retinal waves (the large and long duration ones). As far as we know, this is not a phenotype that has been described, likely because it requires dense recordings across large retinal surface coupled with automatic detection of waves. The phenotype is not surprising: It makes sense that large and long duration waves would account for the majority of the activity that the retina experiences. But accounting for this phenotype is important when investigating the role of genetic and pharmacological manipulations on retinal waves. Generally, one would want to collect every possible retinal wave and ask how a manipulation affects the population average, but if only a minority of retinal waves drive the majority of the activity, research questions should also (and we argue prioritize) ask how a manipulation affects the waves that drive the majority of the activity. The impact of a drug on 10 retinal waves could be hidden if 9/10 waves are small unaffected retinal waves but the 1/10 large, long duration wave that accounted for 80% of the activity is completely prevented by the drug. Thus, the influence of environmental and genetic factors on retinal waves should be reported in two ways: on all retinal waves for full transparency (**figure 2**) and on high-impact waves to get the most accurate picture of how overall retinal activity is affected (**figure 3**).

### Firing patterns of neighboring neurons align with the temporal dynamics of waves

The synchronous activity of nearby RGCs during development is known to degrade around eye opening when retinal waves subside^33,35^. Our results add to the field by demonstrating that from P0 to P14, the STTC of adjacent neurons increases with age. Importantly, the results of our STTC are consistent when we age-match with known data (see P9 data in our results and in Cutts and Eglen, 2014^33^). Thus, we propose an amendment: We predict that STTC values gradually increase during the retinal wave period as the retina develops and precipitously decreases once retinal waves end after P14. The increase in STTC at all distances across the first two postnatal weeks is consistent with the fact that wave frequency is higher during the second postnatal week, meaning that the proportion of times two neurons fire together would be higher in adjacent neurons due to the synchronized nature of retinal waves and in farther neurons due to multiple waves occurring simultaneously.

### Waves in the first postnatal week exhibit better properties for setting up retinotopic maps

A recent study demonstrated the Hebbian mechanism via which the actual propagation of retinal waves is critical for the refinement of retinal axons in the SC^27^. A key factor of retinal waves alluded to in the literature is that as a wave is propagating, neurons just outside of the retinal wave are generally not active, thus allowing for Hebbian plasticity mechanisms to refine exuberant retinal axonal projections. We added a functionality to our toolbox to test the local synchrony of neural activity at the wavefront by tabulating the amount of activity at the wavefront compared to activity just outside of the wavefront. This analysis coupled with STTC allowed us to determine that local synchrony at the wavefront is much higher in the first postnatal week than the second postnatal week, consistent with the fact that the bulk of retinotopic refinement occurs in the first postnatal week^36^. Analysis of local synchrony will be useful for studies that aim to test how various environmental factors or neurodevelopmental disorders affect the key spatiotemporal properties of waves that are important for setting up retinotopic maps.

### Spatiotemporal properties of waves can be dynamic on an hourly basis

Although the spatiotemporal properties of retinal waves have been extensively studied during development^19–22,27,33^, little is known about how waves change throughout the day on an hourly basis. A possible limitation that has led to this lack of information is in the difficulty of segregating waves and analyzing their spatiotemporal properties. Long-term recordings are data-intensive, often consisting of thousands of retinal waves, and cannot efficiently be analyzed without some automation.

Our toolbox resolves this limitation and has allowed us to make a novel discovery regarding waves: Certain spatiotemporal properties of retinal waves can be dynamic or stable in an age-dependent manner. In our recordings, we noticed that wave frequency and wave speed were very stable at their expected age-dependent levels over the course of several hours: For example, wave frequency in the first postnatal week is low compared to the second postnatal week, but these frequencies remained stable over hours for both age groups. By contrast, wave areas exhibited stable properties in the second postnatal week but not the first, with wave areas initially exhibiting a unimodal distribution in the first hour of the recording and exhibiting a bimodal distribution after nine hours.

The dynamic nature of the spatiotemporal properties of waves could perhaps be explained by *ex-vivo* preparations dying over the course of nine hours, but there are several reasons why this explanation is unlikely to be true. First, within the same recording, whereas some spatiotemporal properties can be dynamic (area), others (frequency and speed) are very stable over nine hours, and one would expect the frequency of waves to change drastically if the tissue was dying. Second, the dynamic nature of wave areas is only true in the first postnatal week, just like how the dynamic nature of wave propagation is only true in the second postnatal week. Again, if tissue necrosis was the mechanism of the dynamic nature of the spatiotemporal property, we would expect this to similarly affect retinal waves in the first and second postnatal week. Together, these facts argue against tissue necrosis as an explanation of the dynamic nature of waves.

What then, are possible mechanisms that could cause a spatiotemporal property, like wave area, to be dynamic over the course of hours? One possibility is circadian modulation of retinal activity. Retinal neurons have their own clock proteins^37,38^ and it is possible that excitability of RGCs changes throughout the day, which could lead to broader coverage of the retina at different time periods. Another possible mechanism is that the retina is developing, even in an *ex-vivo* preparation. Future experiments will test which of these mechanisms, if not both, are involved in causing some spatiotemporal properties to be dynamic at specific ages.

The fact that waves are dynamic over the course of hours carries several important implications. One implication, related to methodology and research design, is that future work studying the impact of factors on the spatiotemporal properties of waves needs to be aware that some of these properties can be dynamic throughout the day. The propagation bias of retinal waves has garnered high interest recently^20,34^, and much work is currently being done to determine the exact circuit mechanisms that dictate this bias or whether mouse models of neurodevelopmental disorders exhibit an abnormal propagation bias. Another implication is that since the spatiotemporal properties of waves are critical for the development of the visual system, it now becomes critical to determine the mechanisms that set their dynamic nature. It is possible that certain brain states^39^, via understudied histaminergic and serotonergic brain-to-retina axons^40–42^ or modulation of retinal activity in retinorecipient structures^43^, set the spatiotemporal properties of waves and thus could be critical for the healthy development of the visual system.

### Broad applicability of the toolbox – from pharmacology to other data types

Since the spatiotemporal properties of retinal waves are critical for instructing the development of the visual system, there is a lot of interest in identifying factors that perturb the spatiotemporal properties of waves^20,30,31,44–46^. Unfortunately, tools and methods to study the spatiotemporal properties of waves are specifically designed to work under baseline conditions and fail once a manipulation is done to perturb retinal activity. Our DREADD experiments aimed to demonstrate that WaveMiner retains the ability to segregate and analyze retinal waves even under severe changes in the firing rate of retinal neurons. Thus, WaveMiner enables high-throughput assessment of how retinal waves are perturbed by different pharmacological agents (external factors) or in mouse models of neurodevelopmental disorders. Importantly, the dataset generated from this study or future uses of WaveMiner could be used to train machine learning algorithms to more rapidly detect retinal waves, or to teach us about how retinal waves help train artificial neural networks^47–50^.

Not only is WaveMiner applicable to a wide range of pharmacological experiments, but it can be used to analyze a variety of data types. We have demonstrated that WaveMiner can be used to detect and analyze retinal waves in HD-MEA and calcium imaging data. Our toolbox can be curated to process virtually any type of data containing synchronized events, not just retinal waves, so long as the spatial and temporal information is provided.

## Acknowledgments

This work was supported by NIH grant R00EY030909, Alfred P. Sloan Foundation award FG-2025-24927, and McKnight Scholar award, all awarded to AT. KMO was supported by NIH training grant R25 GM134979 and NSF Graduate Research Fellowship Program. EAG was supported by NIH training grant 5T32GM137793-05.

## Methods

### Ethics

All animal procedures were approved by the Vanderbilt Animal Care and Use Program and conformed to the NIH *Guide for the Care and Use of Laboratory Animals*, the Public Health Service Policy, and the SFN Policy on the Use of Animals in Neuroscience Research.

### Animals

All mice in this study were aged between 0-14 days and of both sexes and were housed in a 12/hr day/night cycle vivarium. Experiments studying the spatiotemporal properties of retinal waves over development as well as long recordings were done using C57BL/6J mice (Jackson stock: 000664). 31 total mice were used in these experiments. Experiments on assessing the spatiotemporal properties of waves following activity manipulations with CNO were done using litters from Vglut2-ires-cre::Gq-DREADD mice (Jackson stocks: 016963 and 026220). 21 total mice were used in these experiments.

### Method Details

#### Retina preparation

Mice were euthanized using an overdose of isoflurane inhalation. Eyes were immediately enucleated and retinas were dissected in ice cold oxygenated under room lights (95% O_2_/ 5% CO_2_) Artificial Cerebrospinal Fluid (ACSF; in mM, 119 NaCl, 2.5 KCl, 1.3 MgCl_2_, 1 K_2_HPO_4_, 26.2 NaHCO_3_, 11 D-glucose, and 2.5 CaCl_2_). We chose to perform all experiments under room light since constant light has no effect on retinal waves across development^12^. Isolated retinas were mounted whole on a MaxWell Biosystems MaxOne chip. The chip was loaded onto a Maxwell Biosystems MaxOne Single-Well High-Density Microelectrode Array (HD-MEA) and the retina was secured using a MaxOne tissue holder. A perfusion system was used to perfuse the retina at a constant rate (2 mL/min) with oxygenated (95% O_2_/ 5% CO_2_) ACSF and maintained at a temperature of 32-34 °C. For all experiments, the ACSF is never recycled.

#### High-Density Microelectrode Array (HD-MEA) recordings

A MaxWell Biosystems MaxOne Recording Unit Hub was connected to the HD-MEA and recorded the retinal activity detected by the electrodes on the MaxOne chip. For all recordings, the electrode configuration consisted of 1024 electrodes (12 reference electrodes, 1012 recording electrodes) distanced 87.5µm apart (MaxLab Live Scope predefined configuration Sparse1x). Following a 1-hour acclimation period, retinal activity was recorded. Only the action potentials were recorded (MaxLab Live Scope setting ‘Record Spikes Only’).

#### Ultra long recordings

9-hour HD-MEA recordings of retinal activity were obtained using the same setup and configurations described above. Due to software limitations, three 3-hour recordings of the same retina were collected consecutively and concatenated post-experiment.

#### Manipulation of retinal waves

To increase overall retinal activity, retinas of VGLUT2Cre::GqDREADD +/+ mice were dissected and loaded onto the HD-MEA. Following a 1-hour acclimation period, a 10-minute recording of retinal activity was collected (referred to as “baseline” or “pre-CNO”). A 10µM solution of the DREADD agonist Clozapine-N-oxide (CNO) in aCSF was then bath-applied to the retina, followed by a 5-minute waiting period, at which point another 10-minute recording was collected (referred to as “+CNO” or “post-CNO”). Note that no control experiments to test for the presence of DREADDs or the ligand CNO are done since the point of this experiment was to artificially raise the floor of neural activity and test whether our toolbox could still segregate waves.

### Data Analysis Details

#### Wave detection algorithm

Hierarchal data (.h5) was uploaded into the wave detection script (written using MATLAB 2024b and 2025b), outputting data organized in a way where every action potential has a time stamp and recorded channel. Combined with information about the spatial location of each channel, we reconstructed movies of retinal activity over time (x by y by t). **At this step, HD-MEA data matches the format of calcium imaging movies (x by y by t), so both types of data can be used moving forward.** To help isolate waves, we first filtered out the tonic firing of each neuron by conducting a burst detection analysis. This algorithm dynamically detects baselines throughout the recording, allowing it to analyze the bursting patterns of neurons and distinguish between tonic and phasic activity. Note that the burst detection analysis was not conducted on the calcium imaging data because it contains calcium transients rather than spiking information. With only phasic activity remaining, the data was fed into the wave detection algorithm. This algorithm scans for active voxels in the xyt movie. Once an active voxel is found, the algorithm determines whether this voxel has active neighboring pixels. If no, the algorithm discards that voxel and scans for the next active voxel. If yes, the algorithm collects spatial and temporal information about all voxels that are connected in space (xy) and time (t), forming a putative wave. Once all connected voxels in space and time are detected, the putative wave is given a unique ID tag, and the participating spatial and temporal voxels are given this ID tag in a resulting wave data matrix. The algorithm then seeks out the next active voxel and the process repeats until all voxels in space and time have been scanned. When the algorithm is done processing the data, the output is a wave data matrix that is the same size as the raw data matrix (xyt).

#### Wave analysis pipeline

The wave data matrices were uploaded into the wave analysis pipeline (written using MATLAB 2024b and 2025b) for analysis of their spatiotemporal properties. This pipeline runs through each recording and analyzes the spatiotemporal properties of each wave in the current recording, compiling the results into a table or structured variable. For a given recording, we calculated the following spatiotemporal properties:

##### Activity impact factor

(AIF) was calculated as:

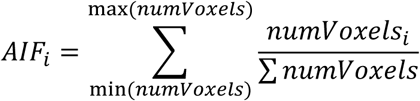

Where *numVoxels* is an array of the number of voxels each wave activates and *i* is the current wave.

The Gini coefficient (*G*) was calculated for each retina as:

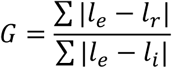

Where *l_e_* is the line of total equality, *l_i_* is the line of total inequality, and *l_r_* is the cumulative AIF curve of the retina. The term in the numerator calculates the area between the line of equality (*l_e_*) and the data (*l_r_*), while the term in the denominator calculates the area between the line of equality (*l_e_*) and inequality (*l_i_*).

##### Inter-wave intervals

(IWI) were calculated for each pixel (*x,y*) as:

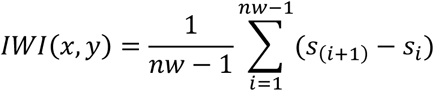

Where *nw* is the number of unique waves that pass through the pixel, *i* is the current wave, and *s* is the start frame of the wave. The IWI for a given recording was obtained by calculating the mean of the pixel IWIs.

Wave frequency was calculated as the reciprocal of IWIs.

Wave area was calculated by isolating the wave into its own matrix, summating the wave in the third dimension (*xyt* to *xy*), counting the number of active pixels, and converting that value to µm² (1 pixel = 87.5 µm).

Wave duration was calculated as:

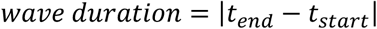

Where *t_end_* is the last frame of activity for a unique wave and *t_start_* is the first frame of activity for a unique wave.

<u>Wave speed</u> was calculated by using a Euclidean distance transform to measure the distances between the leading edge of the wave between pairs of 1-sec time frames. The mean distance traveled by the leading edge was divided by the time between frames (1s) to get the speed of the wavefront in pixels/s, which was then converted to µm/s (1 pixel = 87.5 µm). Top wave speed was obtained by extracting the maximum speed of the wavefront.

<u>Wave initiation sites</u> were calculated by finding the center of mass of the first frame of each wave. Wave initiation bias was calculated as:

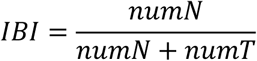

Where *IBI* is the initiation bias index, *numN* is the number of waves that initiated on the nasal half of the retina, and *numT* is the number of waves that initiated on the temporal half of the retina. An *IBI* value closer to 1 indicates a nasal initiation bias, and a value closer to 0 indicates a temporal initiation bias.

Wave vector flow fields were calculated as:

For each wave, the flow of the wave among all participating pixels was calculated for the entire duration of the wave. To calculate the flow of the wave at each pixel, we generated a 3x3 grid of pixels centered on the pixel of interest and calculated the center of mass of the wave within that grid for the entire duration of the wave. The Euclidean distance of each center of mass for that pixel was then tabulated:

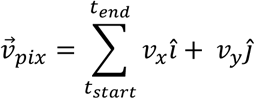

Where, for a given wave, *ν⃗_pix_* is the vector sum over time of a unique participating pixel, *t_start_* is the first frame of the wave, and *t_end_* is the last frame of the wave. *v_x_î* and *v_y_ĵ* are the *x* and *y* vector magnitudes of the difference in the pixel’s grid center of mass between consecutive frames (1s).

The <u>propagation direction</u> of the wave is the average vector of all vectors in the flow field. Wave propagation bias was calculated as:

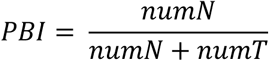

Where *PBI* is the propagation bias index, *numN* is the number of waves that propagate in the nasal direction and *numT* is the number of waves that propagate in the temporal direction. A *PBI* value closer to 1 indicates a nasal propagation bias, while a value closer to 0 indicates a temporal propagation bias. A value of 0.5 indicates no propagation bias.

The spike time tiling coefficient (STTC), as previously published^26^:

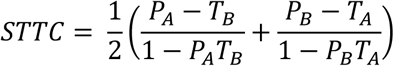

Where *T_A_* is the proportion of total recording time that lies within +/-Δt of any spike from spike train A and *P_A_* is the proportion of spikes from spike train A that lie within +/-Δt of any spike from spike train B. *T_B_* and *P_B_* are calculated similarly.

Local synchrony index (LSI) was calculated as:

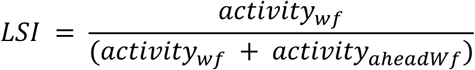

For each wave and frames that are included in the duration of the wave, we tabulated both the amount of activity that is present at the leading wavefront (*activity_nnf_*) and the amount of activity that is present ahead of the wavefront (*activity_a_*_ℎ*ead*W*f*_). For a given frame F_t_ and a subsequent frame *F_t+1_*, we subtracted *F_t+1_* from *F_t_* to obtain the activity ahead of the wavefront.

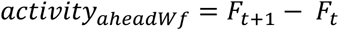

## Data Availability

Processed data is available on our lab’s Github page: https://github.com/TiriacLabCodeShare/waveMiner.

The non-processed dataset will also be uploaded onto that same Github page.

## Code Availability

Analysis code is available on our lab’s Github page: https://github.com/TiriacLabCodeShare/waveMiner.

## Supplemental Information

**Supplemental movie 1.** 3D projection depicting an example wave data matrix, the result of running the wave detection algorithm on retinal wave data. Each color represents a different wave ID.

**Supplemental movie 2.** Example HD-MEA recording before (top, “raw data”) and after (middle, “wave data”) running the wave detection algorithm in a P9 mouse retina. Each color in the wave data represents a different wave ID. The trace on the bottom shows the total number of spikes across the retina over time.

**Supplemental movie 3.** Example HD-MEA recordings before (left, “baseline”) and after (right, “+CNO”) increasing the tonic firing rate of neurons in a P12 Vglut2-ires-cre::Gq-DREADD mouse retina. The original recordings are shown in the top row (“raw data”), and the detected waves are shown in the middle row (“wave data”). Each color in the wave data represents a different wave ID. The traces on the bottom show the total number of spikes across the retina over time.

**Supplemental movie 4.** Example wave detection on a previously published calcium imaging recording of an embryonic mouse retina^12^. The original recording is shown in the top row (“raw data”) and the detected waves are shown in the middle row (“wave data”). Each color in the wave data represents a different wave ID. For this analysis, parameters to detect waves were set to be more conservative to match methods of Voufo et al., 2023^12^, but could be eased to capture more of the small non-propagating events. The trace on the bottom shows the total number of spikes across the retina over time.

---

**Figure S1:**
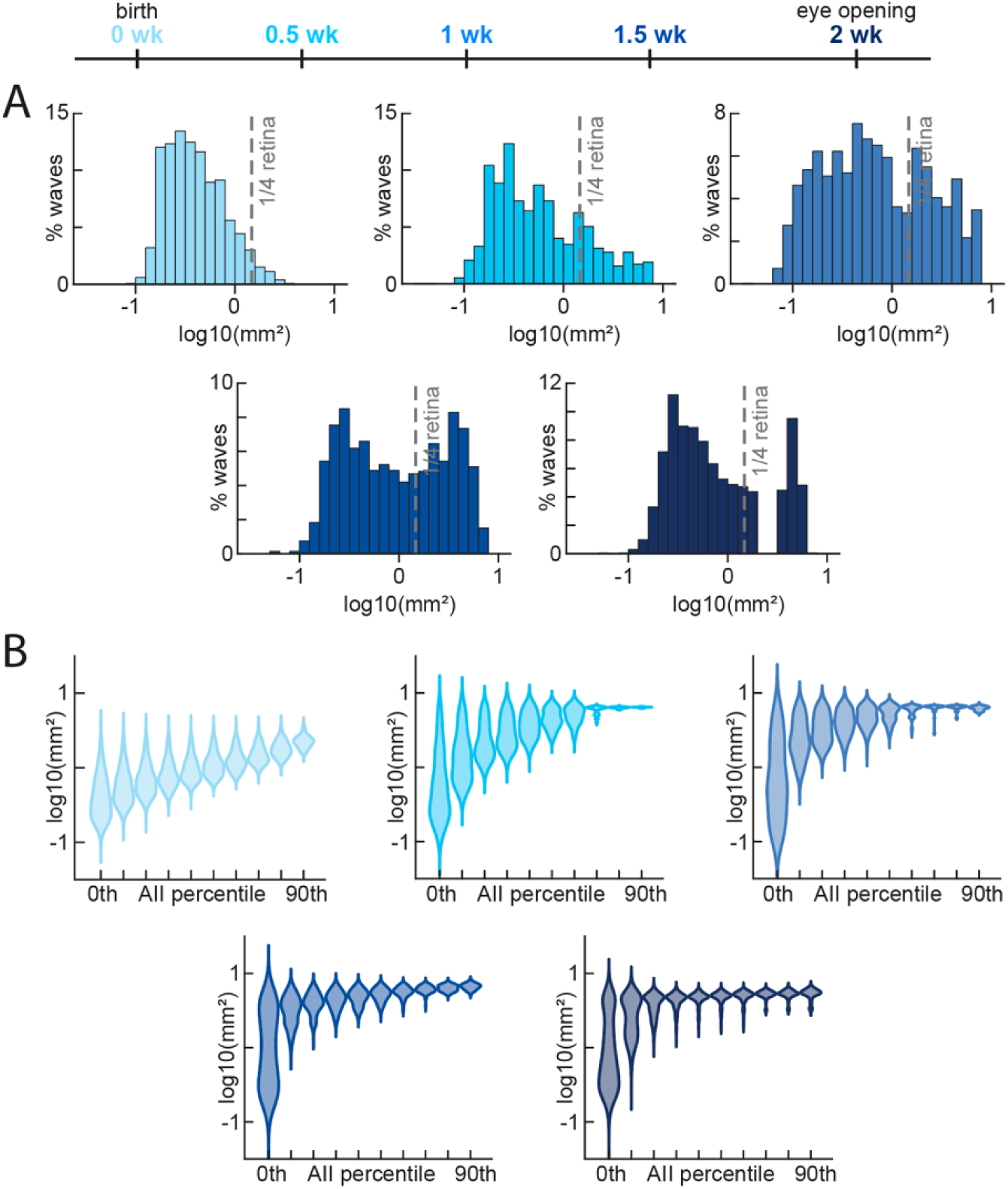
Divergence of wave populations across development. A) Log10 histograms showing the distribution of wave areas across development. B) Log 10 distributions of wave areas within each activity impact factor (AIF) percentile across development.

**Figure S2:**
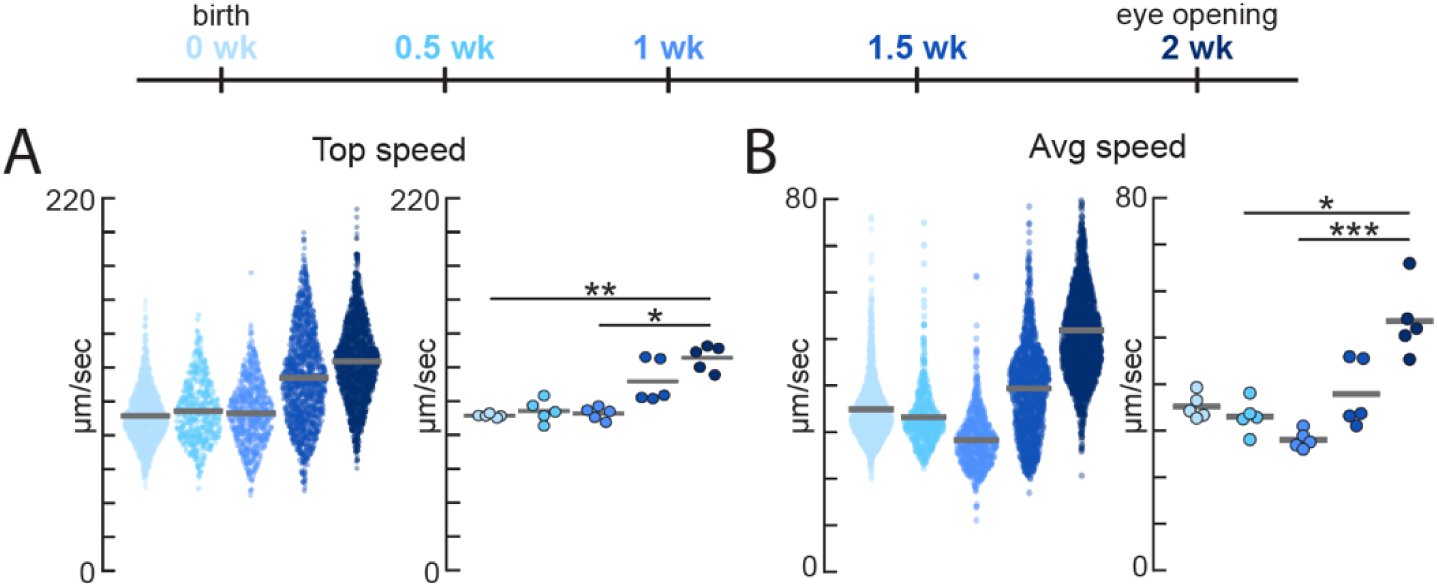
Comparison of average and top wave speeds. A) Top wave speeds across development per wave (left) and per retina (right) of all waves. B) Average wave speeds across development per wave (left) and per retina (right) of all waves.

